# The YjfJ protein is the effector of a second PspA/VIPP1 system that is widespread in the Gammaproteobacteria

**DOI:** 10.64898/2026.11.03.762778

**Authors:** Daniel R. Game, Harry C. Livsey, Marjan W. Van der Woude, Gavin H. Thomas

## Abstract

Maintaining the integrity of the cytoplasmic membrane is essential for bacterial cell survival. Unsurprisingly, prokaryotes have evolved conserved mechanisms by which they sense, maintain, and repair membrane integrity in response to stress. The phage shock protein (Psp) is one such system, understood primarily through work undertaken in the Gammaproteobacterial species *Escherichia coli* and *Yersinia enterocolitica*. The system utilises an effector, PspA, alongside accessory proteins that sense stress and recruit PspA to damaged membranes. Homologues of PspA - part of the PspA/VIPP1 family - are found in a variety of bacterial phyla. Although most work in the Gammaproteobacteria has concerned the canonical Psp response, there have been descriptions of non-canonical PspA/VIPP1-containing systems that warrant further genomic analysis. By analysing the PspA/VIPP1 content of a diverse selection of Gammaproteobacterial genomes, we reveal the existence of two phylogenetically distinct subtypes of PspA/VIPP1 proteins, PspA- and YjfJ-like, which are found at similar frequencies. The YjfJ-like proteins are not only a novel subtype of PspA/VIPP1, having their own unique set of accessory proteins, but in some species, such as the ESKAPE pathogen *Pseudomonas aeruginosa*, comprise the only PspA/VIPP1-containing system. Our study positions the YjfJ-like containing systems on equal footing with the PspA system in terms of their conservation and distribution in the Gammaproteobacteria. This suggests that studies into the biological functions of YjfJ-like proteins would be well-justified from their unique presence in some important human pathogens.

**DATA SUMMARY:** The authors confirm that the supporting data have been provided within the article or in the supplementary material.

**IMPACT STATEMENT:** The discovery of bacterial homologues of mammalian ESCRT-III proteins that remodel membranes stimulated renewed interest in the function of these PspA/VIPP1 family proteins at the bacterial membrane. The first identified bacterial system, discovered over 30 years ago, was the Psp system of the Gammaproteobacterium *Escherichia coli*, and while the regulation of expression of the *psp* genes is well understood, the molecular function of the effector protein PspA at the membrane is still unclear. In this work, using microbial genomics analysis, we find that Gammaproteobacteria contain a second PspA/VIPP1 protein, named YjfJ in *E. coli*, which is widespread and is the only system in the ESKAPE pathogen *Pseudomonas aeruginosa*. Its accessory proteins, which are likely involved in the sensing of stress and activation of the effector, are entirely different to the PspA system which suggests that these are two independent pathways of membrane stress response. In fact, the YjfJ-containing systems appears to be as widespread as the canonical PspA system, suggesting that the study of YjfJ function would shed light on membrane repair systems across the Pseudomonadota (Proteobacteria).

## INTRODUCTION

Bacterial cells are defined by a lipid bilayer that physically separates them from their environment. This cytoplasmic membrane is crucial for the function of cells; it controls entry and exit of metabolites, serves as the site of proton motive force (PMF) generation, and contains other sensory and regulatory functions that allow the cell to sense the external environment (1). The importance of the cytoplasmic membrane to cellular function has led to the evolution of mechanisms to maintain this fragile barrier against a plethora of external physical and chemical threats.

One such system is the phage shock protein (Psp) response, which has been well characterised in selected Gammaproteobacteria. It was discovered through the identification of a 25 kDa protein, PspA, that was synthesised at high levels in *Escherichia coli* following treatment with a number of different membrane-perturbing stresses (2). PspA is a PspA/VIPP1 family protein (InterPro entry IPR007157) and is thought to protect the inner membrane and maintain PMF in response to membrane damage (1,2). PspA/VIPP1s sit within the ESCRT-III superfamily of membrane remodelling proteins that are found across all three domains of life (3). While PspA is the effector in the Psp response, the system contains additional accessory proteins encoded within the same operon, the most well-characterised of which are PspB and PspC (4–6). These two proteins are located in the inner membrane and are involved in sensing stress and recruiting PspA to the damaged membrane (7). Research has also indicated that the function of this membrane stress response is linked to virulence in some key Gammaproteobacterial pathogens; for example, in *Yersinia enterocolitica* the *psp* operon is required for virulence, and in *Salmonella enterica* serovar Typhimurium it is needed for resistance to host-derived CAMPs and pathogenicity (8–10).

To date, *in silico* studies of PspA/VIPP1 systems have taken a broad view across all prokaryotes and archaea and have revealed the diversity of these proteins and their accompanying accessory proteins (11,12). The *psp* operon as it is found in the Gammaproteobacteria is restricted to only a few phyla outside of the wider Proteobacteria (11). Instead, other arrangements of PspA/VIPP1 encoding gene clusters are more prevalent across all prokaryotes and the accessory proteins can vary substantially from those in the *psp* operon (11,12), suggesting that bacteria have found multiple ways to activate the same or similar effector proteins. PspA/VIPP1-containing systems containing different accessory proteins to the canonical *psp* operon have been studied in the Gammaproteobacteria. The biofilm-associated cluster (*bac*) found in *Pseudomonas aeruginosa* is one example, with the PspA/VIPP1 protein BacB implicated in the formation of biofilms (13). The accessory gene, *bacA*, is a chaperone thought to act as a post-transcriptional regulator of operon expression (14). Notably *E. coli*, the organism in which the *psp* operon was first described, also has a second PspA/VIPP1-containing system (15,16). The *yjfIJKL* cluster was found to be upregulated in strains evolved in the laboratory to grow at higher temperatures than the wild-type (16), but is poorly studied and the role of the accessory proteins in this gene cluster are unknown.

A more focused genome analysis of PspA/VIPP1s and their gene clusters can reveal nuances and trends hidden by the lack of granularity offered by broader analyses. To do this we performed in-depth analysis on a sample of Gammaproteobacterial genomes that were chosen to be representative of the Class’ diversity. Firstly, we analysed the distribution of PspA/VIPP1s across the representative genomes, before analysing the phylogeny of the protein family, and determining the genome contexts of each PspA/VIPP1. In doing so we have determined that PspA/VIPP1s are not ubiquitous in the Gammaproteobacteria, and that they have been lost at multiple points in evolution. Our analysis has also revealed that two phylogenetically distinct subtypes of PspA/VIPP1 can be found across the Gammaproteobacteria, and that protein subtype correlates directly with genome context.

## METHODS

### Selection of Gammaproteobacterial genomes

Analysis was performed in March 2024, and at that point there were 20,345 fully assembled genomes from the Gammaproteobacteria available on the National Centre for Biotechnology Information (NCBI) database. Our representative dataset contains 84 genomes that reflect the diversity of the Orders within the Gammaproteobacteria. The selection was based on a previously reported phylogeny (17), which was adapted to reduce the overrepresentation of some genera and include some more recently discovered Orders, reducing bias. The Betaproteobacterium *Burkholderia cepacia* was included as a PspA-containing outgroup. All genomes in the dataset are included in Table S1.

### Identifying PspA/VIPP1 proteins

Searches were performed against the InterPro database using the family IPR007157 (PspA/VIPP1). Hits were cross-checked against the genome of the relevant strain in NCBI. Genomes with no hits from the InterPro analysis then had their proteins analysed manually using BLASTp (with default parameters), using *E. coli* PspA and YjfJ (UniProt IDs P0AFM6 and P0AF78 respectively) as query sequences to confirm that there were no significant matches (E value < 0.01).

### Phylogenetic analyses

Both 16S rRNA and PspA/VIPP1 phylogenetic trees were generated using the same workflow, first sequences were aligned using the MAFFT (v7.526) alignment programme (18,19). FastTree 2.2 was then used to infer approximately-maximum-likelihood phylogenetic trees based on the alignment (20). Pseudogenes were excluded from the phylogeny to avoid distorting the tree with poor alignments. The resulting phylogenetic trees were visualised and edited using the iTOL software (v7) (21). *B. cepacia* 16S rRNA or PspA was used as the outgroup as relevant for phylogenetic trees.

### Gene cluster analysis

Gene clusters were defined according to criteria used previously, as having the same strand orientation and with an intergenic distance no longer than 200 bp (22–24). Divergently transcribed genes were included in the cluster analysis to reflect the role of PspF in the regulation of the Psp system (25,26). Gene cluster analysis was performed manually using the relevant genome from NCBI. InterProScan was used to predict membership of protein families and domains from the sequence (27,28).

## RESULTS

### PspA/VIPP1s are not ubiquitous in the Gammaproteobacteria

To identify the presence of PspA/VIPP1 sequences in our dataset, species were first checked against the PspA/VIPP1 database in InterPro, verified by manual checks with *E. coli* PspA and YjfJ. Across the 84 genomes, we found that PspA/VIPP1 proteins were not evenly distributed (Fig. 1). Just over half (48 of 84 species) were found to contain a PspA/VIPP1 protein, illustrating that these proteins are not universally present in the Gammaproteobacteria. Species containing at least one PspA/VIPP1s were readily found in the Enterobacterales, Vibrionales, Alteromonadales, and the Pseudomonadales, with other Orders containing more sporadic distributions.

**Figure 1.**
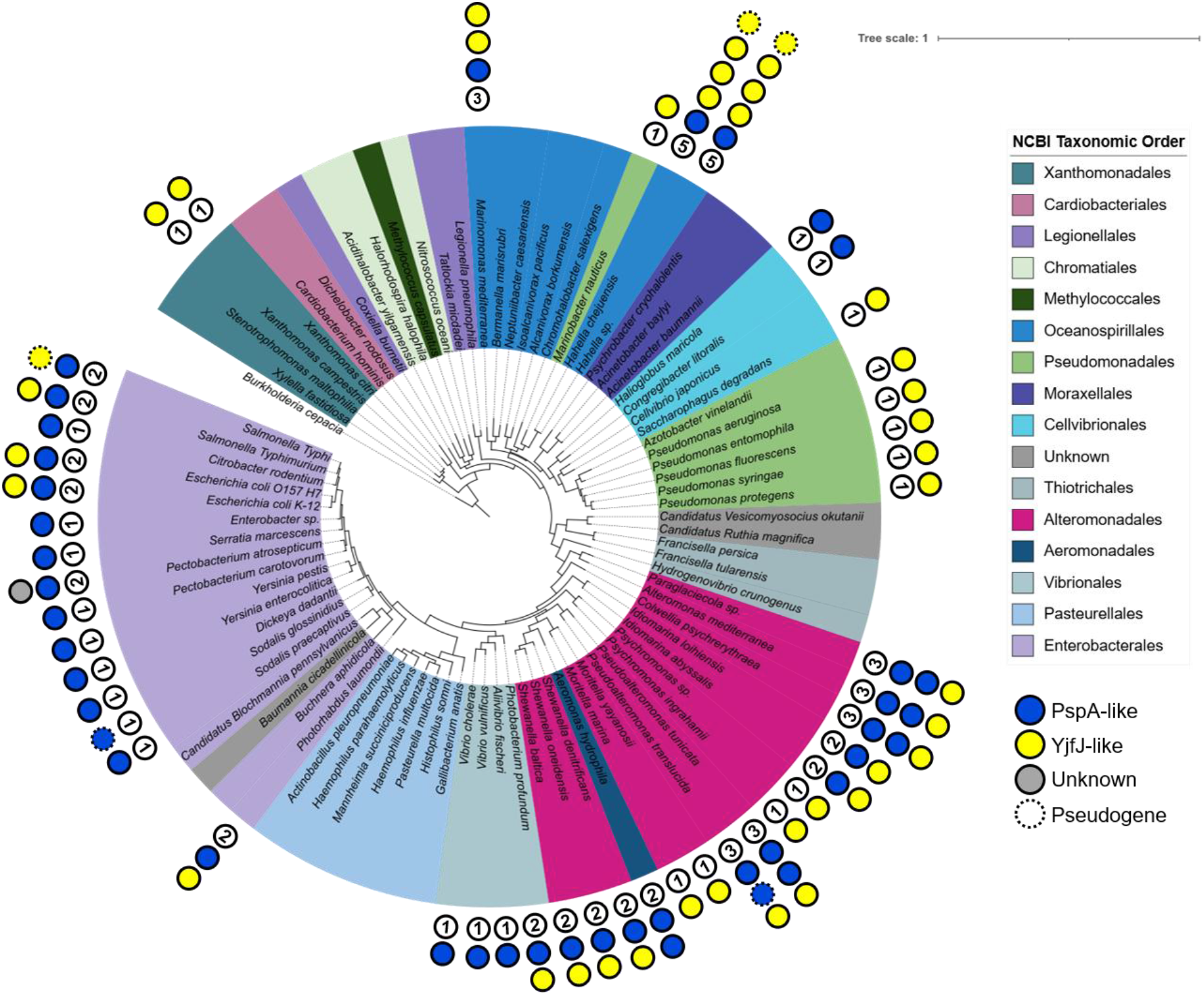
PspA/VIPP1s are not ubiquitous across the Gammaproteobacteria. An approximate-maximum-likelihood phylogenetic tree of 16S rRNA for each species in the dataset, rooted to the Betaproteobacteria B. cepacia. Species are coloured by their taxonomic Order according to NCBI Taxonomy (as of 12 March 2024). The inner circle represents the number of PspA/VIPP1 sequences present in each species. Each circle represents a PspA/VIPP1 with the colour showing subtype – PspA-(blue) or YjfJ-like (yellow), or unknown (grey). Circles with a dashed outline denote pseudogenes.

There are clear groups within the tree that lack any PspA/VIPP1 system, which is consistent with previous broad analyses of Psp domains (12) (Fig 1.). For example, the Pasteurellales and Moraxellales, which contain human pathogens including *Haemophilus influenzae* and *Acinetobacter baumanii*, respectively, contain no PspA/VIPP1s. This absence was confirmed to be consistent across the whole Order using BLASTp with *E. coli* PspA and YjfJ (data not shown). As the PspA/VIPP1s proteins share similarity to eukaryotic ESCRT-III-like proteins and are found across all three domains of life, they are thought to have been present in the last universal common ancestor (LUCA). Therefore, these proteins were likely retained at the root of the Gammaproteobacteria tree (3). Our analysis suggests selective gene loss in particular Orders, and as the Orders lacking any PspA/VIPP1 proteins are polyphyletic this suggests that loss of these genes has occurred on multiple evolutionary occasions. An example of more recent selective loss can be seen within the two *Sodalis* species. The free-living bacterium *S. praecaptivus* has an intact PspA/VIPP1-coding gene, while its symbiotic relative *S. glossinidus* has inactivated its copy. Evolving to life in the tsetse, *S. glossinidus* has pseudogenised both *pspA* and *pspB,* showing that their function is non-essential in a more environmentally stable niche (29,30).

Examining species that do contain functional systems, we can observe that the well-studied Enterobacteriales contain one or sometimes two systems, while other Orders such as the Alteromonadales can contain three. Remarkably, a number of bacteria in the Oceanospirallales contain five systems (Fig. 1).

We considered the possibility that the number of PspA/VIPP1s was simply a result of genome size (Fig. S1). We found that genomes containing PspA/VIPP1s were significantly larger than those without. This indicates that larger genomes are more likely to have PspA/VIPP1s; however, there are exceptions such as *Acinetobacter baumanii*, that have medium sized genomes (∼ 4 Mb) but still lack any PspA/VIPP1 systems (Fig. 1). Nevertheless, when we exclude genomes with no hits and look at any correlation between genome size and the presence of one or more PspA/VIPP1s we see there is no statistically significant relationship. This seems to indicate that something other than large genomes and the acquisition of genes is driving the selection for multiple PspA/VIPP1s within a species.

### PspA and YjfJ are phylogenetically distinct subtypes of PspA/VIPP1

All PspA/VIPP1 sequences in the dataset were used to infer a phylogenetic tree of PspA/VIPP1 diversity across the Gammaproteobacteria (Fig. 2). Surprisingly, the sequences fell into two distinct and separate clades – one containing *E. coli* PspA and the other containing *E. coli* YjfJ, with a third deeply branching group containing a small number of more distant members of the family (Fig. 2).

**Figure 2.**
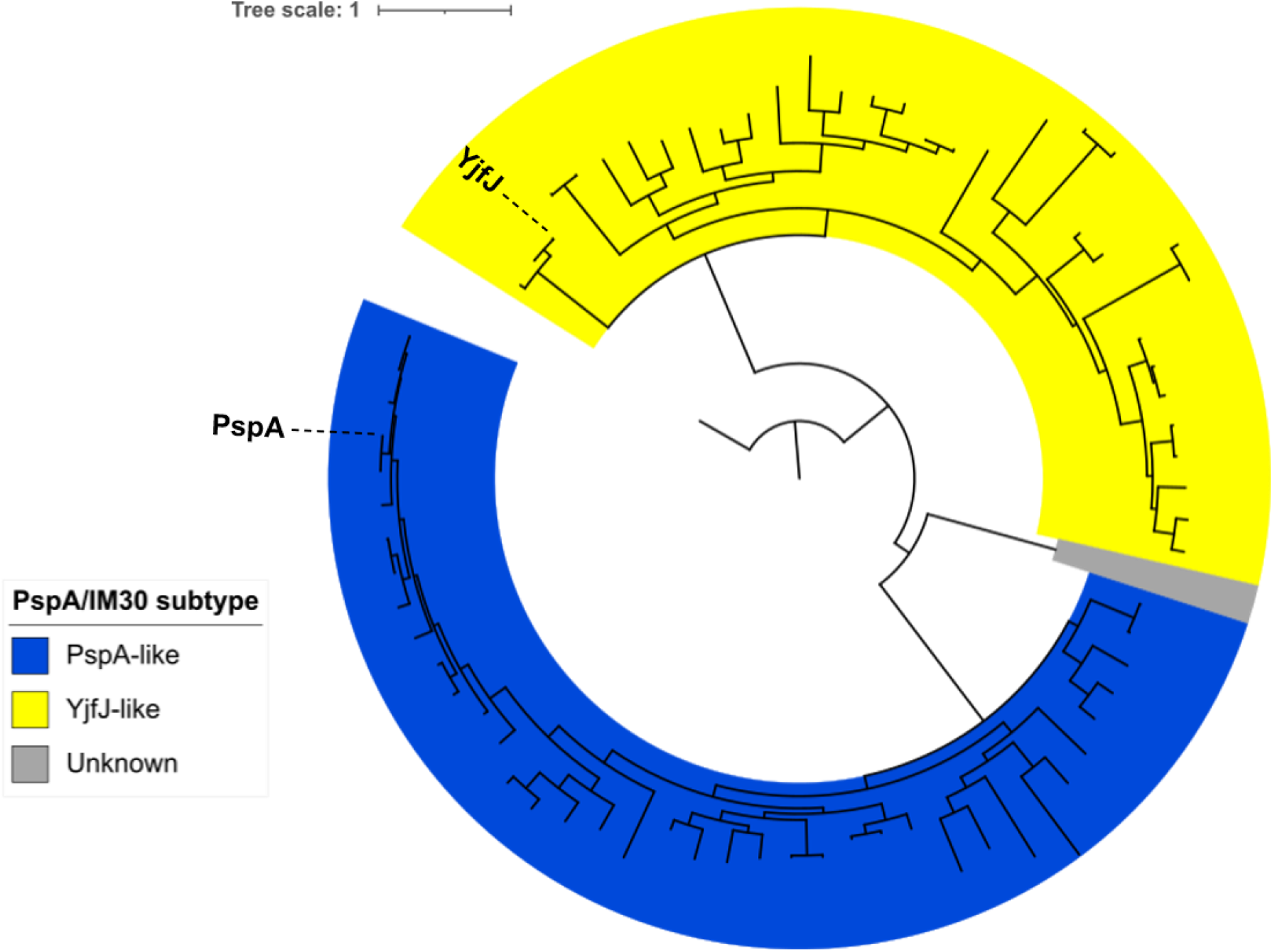
PspA and YjfJ are phylogenetically distinct subtypes of PspA/VIPP1. An approximate-maximum-likelihood phylogenetic tree of all PspA/VIPP1 sequences in our dataset, excluding pseudogenes, rooted to B. cepacia PspA. Clades are coloured by the presence of E. coli PspA (blue), YjfJ (yellow), or unknown (grey). The location of E. coli YjfJ and PspA are labelled.

The distinction between the two *E. coli* PspA/VIPP1s has been reported previously (15), but this result shows that they are both found widely across the Gammaproteobacteria, in approximately equal frequency. Expanding this analysis to all PspA/VIPP1s from the Gammaproteobacteria listed in InterPro (2491 sequences) confirms the pattern observed with the selected genomes and supports the idea that these two proteins, PspA and YjfJ-like, are phylogenetically distinct and can be classified as distinct subtypes of PspA/VIPP1 (Fig. S2).

Having established these two groups of PspA/VIPP1 proteins, we examined our selected genomes to highlight some patterns of distribution of systems across the Gammaproteobacteria (Fig 1). While the Enterobacterales have predominantly PspA-type systems (55 out of 573), reflecting the discovery of PspA in *E. coli*, there is a striking difference in the Pseudomonadales that almost exclusively have YjfJ-like systems (247 out of 265) (Fig. 1, Fig S3). This includes the important ESKAPE pathogen *Pseudomonas aeruginosa* and other well-studied Pseudomonads such as *P. syringae* and *P. fluorescens* (Fig. 1). Across other Orders the patterns are less biased to one or the other, for example the Alteromonadales have a relatively similar spread of both systems (Fig. 1, Fig. S3).

In many cases there are also variations within an Order, clearly seen in the Enterobacterales, where the YjfJ-like systems are only found in one branch of the tree that contains *E. coli* and *S.* Typhimurium, but is absent in the rest and even appears to have been lost relatively recently in the pathogenic *E. coli* relative *Citrobacter rodentium* (Fig. 1). Intriguingly the human host-restricted relative of *S.* Typhimurium, *Salmonella enterica* serovar Typhi, has pseudogenised its YjfJ-like PspA/VIPP1 (Fig. 1).

We also note that the *Hahella* species in our analysis appear to have 4, and at one time possibly 5, different PspA/VIPP1 proteins. Our data shows this is through expansion of the YjfJ-like systems, which are the most in a single organism we are aware of, although the physiological reasons for this adaptation in this marine bacterium are not apparent.

The unknown subtype in the in-depth tree (Fig. S2, grey) highlighted a third grouping of more distantly related proteins. One of these is highlighted in our analysis in Fig. 1 from *Pectobacterium atrosepticum* (Protein ECA0777, UniProtID Q6D942), which is absent in its close cousin *P. carotovorum* (Fig 2). This lack of conservation within the Genus – coupled with the comparative uniqueness of the sequence – is indicative that the second sequence has been acquired by horizontal gene transfer. BLASTp analysis of ECA0777 supports this hypothesis with a match from the closely related species *P. braziliense*, followed immediately by a series of homologues from Betaproteobacteria and Firmicutes.

### PspA- and YjfJ-like PspA/VIPP1s have distinct genetic environments

Accessory proteins are understood to be key to the function of PspA/VIPP1s in bacteria and are encoded in gene clusters with the PspA/VIPP1 proteins themselves (4,13,14,31,32). Given that our analysis has indicated that PspA- and YjfJ-like sequences are distinct subtypes of PspA/VIPP1, we wish to examine their genome context to see if they also are linked to a distinct sets of genes encoding accessory proteins, suggesting different physiological functions for the two systems.

We used methods previously employed for computational operon prediction in bacteria (22–24), considering genes to be part of a cluster if they are transcribed in the same strand and have an intergenic distance is < 200 bp from their neighbouring up or downstream genes, with the PspA/VIPP1 encoding gene as the starting point for each cluster. In addition, divergently transcribed genes from the primary operon were included, given that the transcriptional activator for the *psp* operon, *pspF*, is divergently transcribed from the rest of the operon (25,33) and that this is a common layout for co-regulated genes in bacteria (34,35).

The patterns of gene clustering around the PspA/VIPP1-encoding genes are highly conserved within the *pspA*-containing clusters (Fig 3A), while there is more variation in the *yjfJ*-containing clusters (Fig. 3B) with some clear patterns of conserved linked genes. However, the two systems share no common genes beyond the one encoding the PspA/VIPP1 protein, which strongly supports the idea that these encode entirely different stress response systems, likely sensing different membrane-based signals of damage.

**Figure 3.**
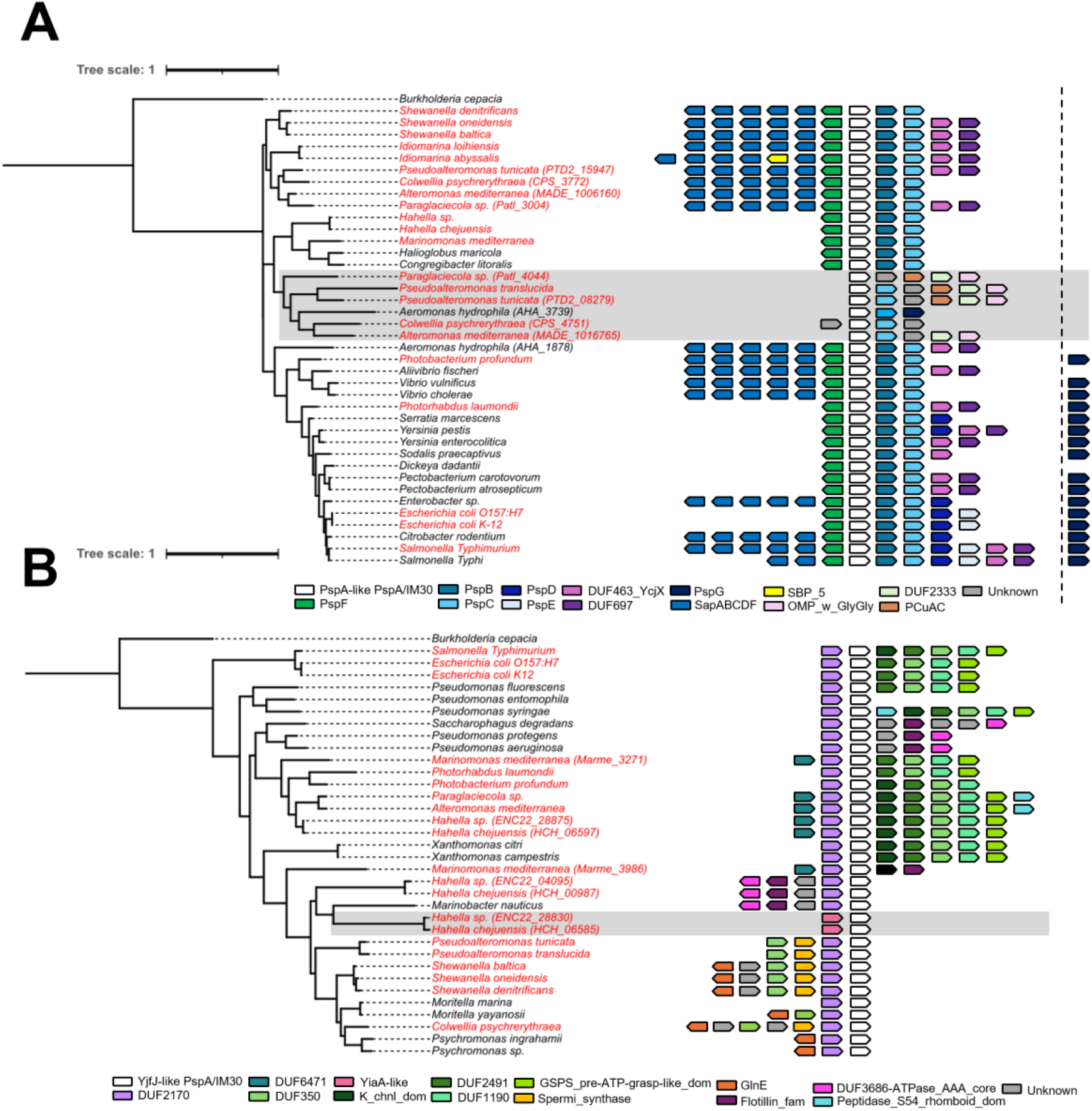
PspA- and YjfJ-like PspA/VIPP1s have distinct genetic environments. Gene clusters for each PspA/VIPP1 shown alongside approximate-maximum-likelihood phylogenetic trees for both PspA-(A) and YjfJ-like (B) PspA/VIPP1s. Gene clusters were defined as having the same strand orientation and an intergenic distance < 200 bp with the PspA/VIPP1 as a central point. Divergently transcribed genes were included to reflect the role of PspF in the regulation of the Psp system. Colours indicate protein identity as determined from sequence using InterProScan (genes not to scale). When multiple IM30s are present in an individual genome the specific locus tag is indicated; species labelled in red have both PspA- and YjfJ-like proteins. Grey shading shows non-canonical PspA/VIPP1 gene clusters for PspA- and YjfJ-like proteins. Genes appearing in two or more clusters are shown, additional genes meeting the definition for membership of a gene cluster are listed in Table S2 and all InterProScan results can be found in Table S3.

### The *pspFABC* genes form the core component of the Gammaproteobacterial Psp system

The *psp* genes as initially described in *E. coli* comprise seven genes, namely *pspF*, *pspABCDE*, and *pspG*, organised in three transcriptional units (26,33,36). Examining the genes associated with *pspA* across the Gammaproteobacteria, we see *pspF* and *pspABC* being widely conserved, with *pspDE* being less conserved and additional genes seen in some cases. While it has been suggested that the core elements of the Psp response are PspA and PspC (12), this analysis supports a core role of the four genes *pspFABC* in the Gammaproteobacteria.

Exceptions to this that lack *pspB* are observed in some bacteria, such as *Colwellia psychrerythraea*, where the PspA-like encoding gene *CPS_4751* lies directly upstream of a PspC domain-containing gene and downstream of a gene of unknown function. However, this is the second PspA-like system in this bacterium (grey box in Fig. 3A shows ‘second’ PspA-like-containing systems) in addition to a canonical *pspFABC* containing the PspA encoding gene *CPS_3772*.

We have established *pspFABC* as the core genes of the Psp system, but the downstream genes vary in different species (Fig. 3A). In *E. coli* the downstream genes are *pspDE*, but in *Y. enterocolitica ycjXF* are present which are found in a number of PspA-like contexts throughout the Gammaproteobacteria. Interestingly, *S.* Typhimurium and *S.* Typhi have both *pspDE* and *ycjXF* downstream of *pspFABC*. Transcriptomics data from the SalComD23580 database (37) shows that *pspABCDE-ycjXF* and *pspG* are all upregulated in a macrophage infection context (Fig. S4). This indicates that these genes are all co-regulated in *S.* Typhimurium and that both pairs of accessory genes can function together within a PspA-containing system.

Unusually, *pspE* appears to be upregulated in additional conditions independently of the rest of the operon (Fig. S4). This suggests an additional promoter specific for the sole transcription of *pspE* which would be independent of PspF as the transcription activator. This idea is supported by the iModulonDB for both *Salmonella* (CORE dataset) and *E. coli* (PRECISE815) which groups *pspABCD* and *pspG* into the same PspF iModulon of co-regulated genes, but excludes *pspE* (38,39). Interestingly, Fig. 3A shows that *pspE* is only conserved in three species in our dataset - *E. coli*, *S.* Typhimurium, and *C. rodentium*. This may provide clues as to the role of PspE in the Psp response, and what necessitates its own separate promoter.

Although not directly in the *psp* locus, *pspG* is also regulated by PspF (36,40). The genomes that contain *pspG* are restricted to the Enterobacterales and the Vibrionales (Fig. 3A). Interestingly, *A. hydrophila* does have a *pspG* but unusually it is downstream of its second PspA-like PspA/VIPP1 and a second *pspC*. The function of PspG, like some of these other accessory proteins, is not clear, but it is not a quirk of *E. coli* alone.

### Co-localisation of the *sap* genes with the *psp* operon, suggests possible integration of cell membrane damage reduction systems

In addition to defining the core *pspFABC* gene set, analysis of the flanking genes found a common association with another set of 5 genes, often encoded directly downstream from *pspF* (Fig. 3A). This linkage is not seen in *E. coli*, where there are additional genes interposed between them, but it appears common in other pathogenic genera such as *Salmonella* spp. and overall is seen in 18 out of 33 canonical PspA-like arrangements.

The linked genes encode the Sap ABC transporter system, which has been implicated in resistance to cationic antimicrobial peptides (CAMPs) (41,42), which often perturb the function of the cytoplasmic membrane. Both systems are implicated in protecting the membrane from damage which supports the connected functions of the Psp and Sap systems, and previous reports have shown that ∼87% of *sap* operons have the Psp system upstream (43). These joint sets of genes have also been seen to have been mobilised and amplified together in antibiotic resistance plasmids found in a number of clinical strains of different Enterobacteriaceae, including *S.* Derby (44) and the ESKAPE pathogens *Enterobacter* spp. (45–47), suggesting some conserved function in antibiotic resistance. We sought evidence for possible coregulation of these genes under different stress conditions, but there was no overlap in the regulation of the *sap* and *psp* genes in SalCom (Fig. S4) or iModulonDB for *Salmonella* or *E. coli*.

### The *yjfJ*-containing clusters all encode a DUF2170 domain-containing protein directly upstream of the PspA/VIPP1 effector

The most striking observation from the *yjfJ*-containing clusters (Fig. 3B) is the association of the PspA/VIPP1 encoding gene with an upstream gene encoding a DUF2170 protein, a domain that has been linked to protein chaperone activity. The existence of a conserved two gene chaperone-PspA/VIPP1 dyad has been described before (11,14), but here we can clearly see that this connection is specifically to the YjfJ-like systems and not the PspA system.

The only exceptions to this DUF2170-YjfJ-like PspA/VIPP1 association are where a Species has more than one YjfJ-like system. At least one of the genetic environments will contain a DUF2170 protein, for example in *H. chejuensis* where *HCH_06585* lacks the chaperone, but both other *yjfJ*-containing environments do have copies of this gene (Fig. 3B). Even when the genetic environment encodes different accessory proteins, as is shown in Fig. 3B and described by Ravi *et al.* (11), the DUF2170-YjfJ-like PspA/VIPP1 remains conserved. Interestingly, this genetic association between DUF2170 and PspA/VIPP1 proteins has been shown to be conserved beyond the Gammaproteobacteria. Ravi *et al* (11) have shown that examples of this association are found in phyla as diverse as the Alphaproteobacteria, Firmicutes, FCB superphylum, and Bacteroidetes.

## DISCUSSION

The successful maintenance of the function of the cytoplasmic membrane is important for cell viability, and the PspA/VIPP1 proteins are involved in protecting this membrane through mechanisms that are not fully understood. For the *E. coli* Psp system, PspA is likely the effector protein working on the membrane, with the accessory proteins appearing to aid this process by sensing the signal of damage in the membrane and/or helping the PspA effector to function, although again details of direct mechanisms are not entirely clear. Our analysis has clearly identified a second PspA/VIPP1 system present in *E. coli* but also widespread across the Gammaproteobacteria. The function of this system, with a YjfJ-like effector, is even less well understood.

The occurrence of these two primary systems follows no clear pattern across the Gammaproteobacteria. This is nicely illustrated by their occurrence in the four Gammaproteobacterial ESKAPE pathogens, with *Enterobacter* spp. and *Klebsiella pneumoniae* having the canonical Psp system, *P. aeruginosa* having a YjfJ-containing cluster, and *A. baumanii* having no PspA/VIPP1 proteins. There is also no strong correlation between having one system and having the other, suggesting that different evolutionary selection is operating on both systems and that they have discrete physiological functions, consistent with their unique sets of accessory genes. The fact that *A. baumanii* and other Gammaproteobacteria lack any systems also illustrates their non-essential nature, unless this is compensated by an independent stress response in these organisms.

The Psp system, encoded by *pspFABC*, appears to be limited to Proteobacteria, with PspB being most restricted to this phylum (11). This is similar to other Phylum-specific accessory proteins in PspA/VIPP1 systems, such as PspM and PspN which are accessory proteins found in the *Mycobacterium tuberculosis* and other Actinobacteria (11). In contrast to the Psp system, the distribution of the DUF2170-PspA/VIPP1 pair is not restricted to a single bacterial Phylum. Ravi *et al*. (11) refer to the DUF2170 domain as a CesT/Tir-like chaperone protein, reflecting its structural similarity to type-III secretion system chaperones (14,48). One could speculate that the chaperone is involved in the folding/assembly of the PspA/VIPP1 protein, most likely by binding to monomers and keeping them in a partially unfolded state and inhibiting functionality. The consistent localisation of the chaperone gene directly upstream of the effector gene is perhaps consistent with this idea, such that the chaperone is always made before the effector.

Clearly the function of the YjfJ-like systems needs experimental study and the best clues currently come from work in *P. aeruginosa* where the protein has been designated BacB. Loss of this gene results in a number of measurable phenotypes, many of which can be explained by the reduction in level of rhamnolipid production (14). The DUF2170 protein, BacA, has been proposed to be a negative regulator of expression of the other *bac* genes in the cluster (14). Wallart *et al.* (14) draw a parallel to the ExsC protein which is involved in regulating the Exs Type-III secretion system, but through protein-protein interactions and not through direct transcriptional regulation Hence, it seems likely that the YjfI-like/BacA/DUF2170 protein may bind YjfJ/BacB in some manner and this could represent a resting inactive complex that provides a pool of PspA/VIPP1 that could be recruited directly to the membrane. The function of the chaperone may end here, or it might bind another partner and hence transduce some regulatory signal of system activation. Information on the biochemical and functional role of the *E. coli* Yjf system is lacking, with the only phenotypes suggesting a role in growth at elevated temperature (16,49). The suggestion that each system has distinct physiological roles is also supported by the work of Seo *et al*. (50), who reproduced secretin-based toxicity in *Pseudomonas aeruginosa*, for which PspA plays a role in both *E. coli* and *S.* Typhimurium and while identifying various *P. aeruginosa* genes that protected from this stress, YjfJ (BacB, PA3731) was found to have no role in this process.

The direct study of these YjfJ-containing systems is very limited, however our work has shown that they are a unique, well-conserved subtype of PspA/VIPP1 that is present as frequently as PspA in the Gammaproteobacteria. Given our understanding of the role PspA can play in bacterial virulence, and the conservation of YjfJ-like PspA/VIPP1s in important bacterial pathogens, studies into the biological function of these systems is well-justified.

## ACKNOWLEDGEMENTS

The authors would like to thank Vytaute Jurgeleviciute, Reyme Herman, and Benjamin Willson for their constructive feedback on the manuscript.

## SUPPLEMENTARY FIGURES

**Figure S1.**
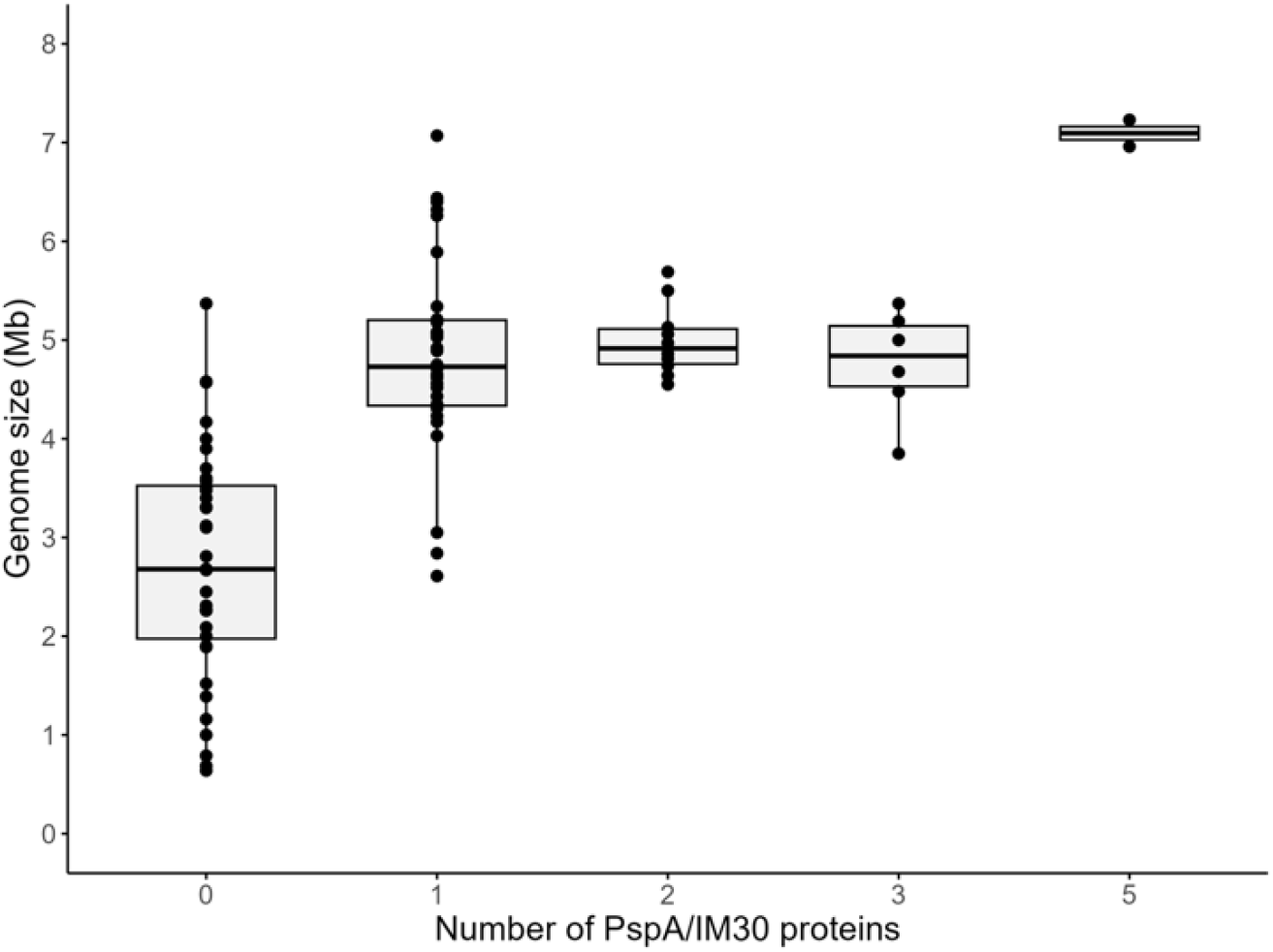
Box plot of genome size (Mb) vs the number of PspA/VIPP1 proteins per species for our dataset. The largest genome size that contains no PspA/VIPP1 is Azotobacter vinelandii (5.37 Mb). The smallest genome with at least one PspA/VIPP1 is Idiomarina abyssalis (2.61 Mb). Genomes containing PspA/VIPP1s were significantly larger than those without (Wilcoxon rank-sum test W = 115, p = 1 x10-11; Median size: 2.68 Mb vs 4.88 Mb). There is no correlation between one or more PspA/VIPP1s and genome size (Spearman’s rank correlation coefficient ρ = 0.20, p = 0.17).

**Figure S2.**
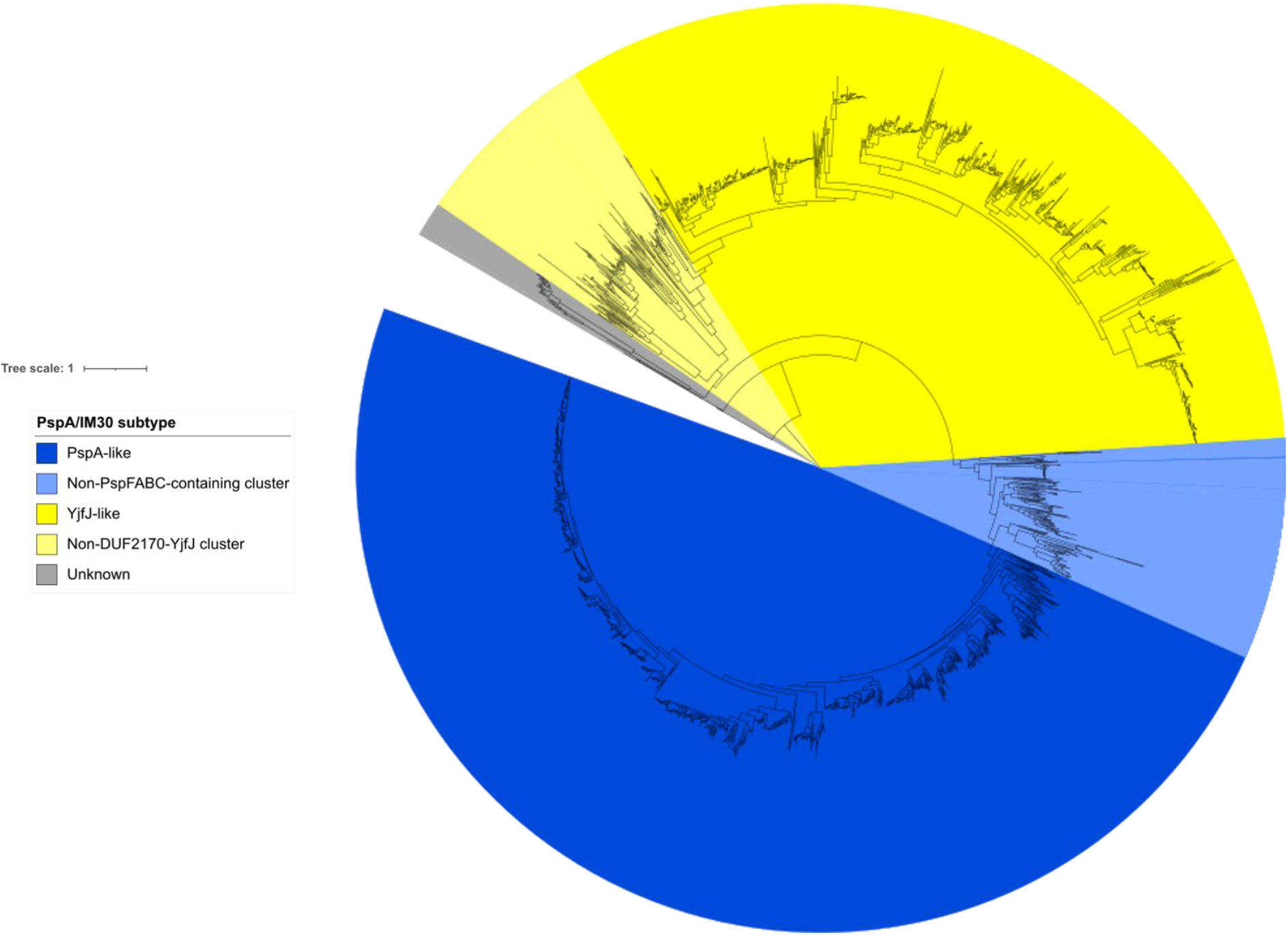
An approximate-maximum-likelihood phylogenetic tree of all Gammaproteobacterial PspA/VIPP1 sequences in the InterPro family IPR007157 (2419 sequences) rooted to B. cepacia PspA. Clades are coloured by the location of E. coli PspA (blue) or YjfJ (yellow), or an unknown subtype (grey). PspA-like proteins not found in a pspFABC genome context are marked in light blue; YjfJ-like proteins not found alongside DUF2170 chaperones are coloured light yellow.

**Figure S3.**
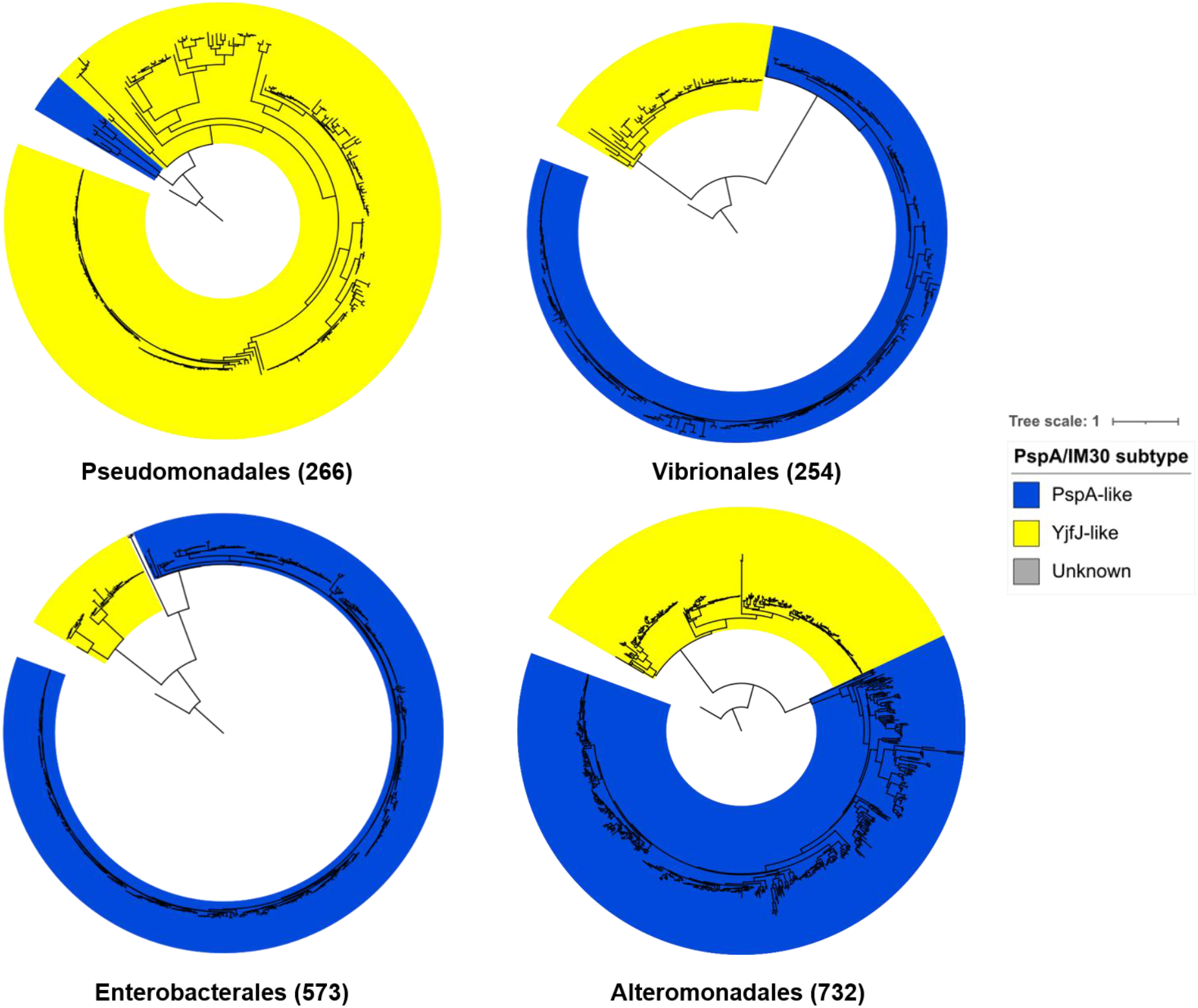
Individual approximate-maximum-likelihood trees for the Pseudomonadales, Vibrionales, Enterobacterales, and Alteromonadales Orders of the Gammaproteobacteria, rooted to B. cepacia PspA. Brackets indicate the number of proteins in each phylogenetic tree. Clades are coloured by the location of E. coli PspA (blue) or YjfJ (yellow).

**Figure S4.**
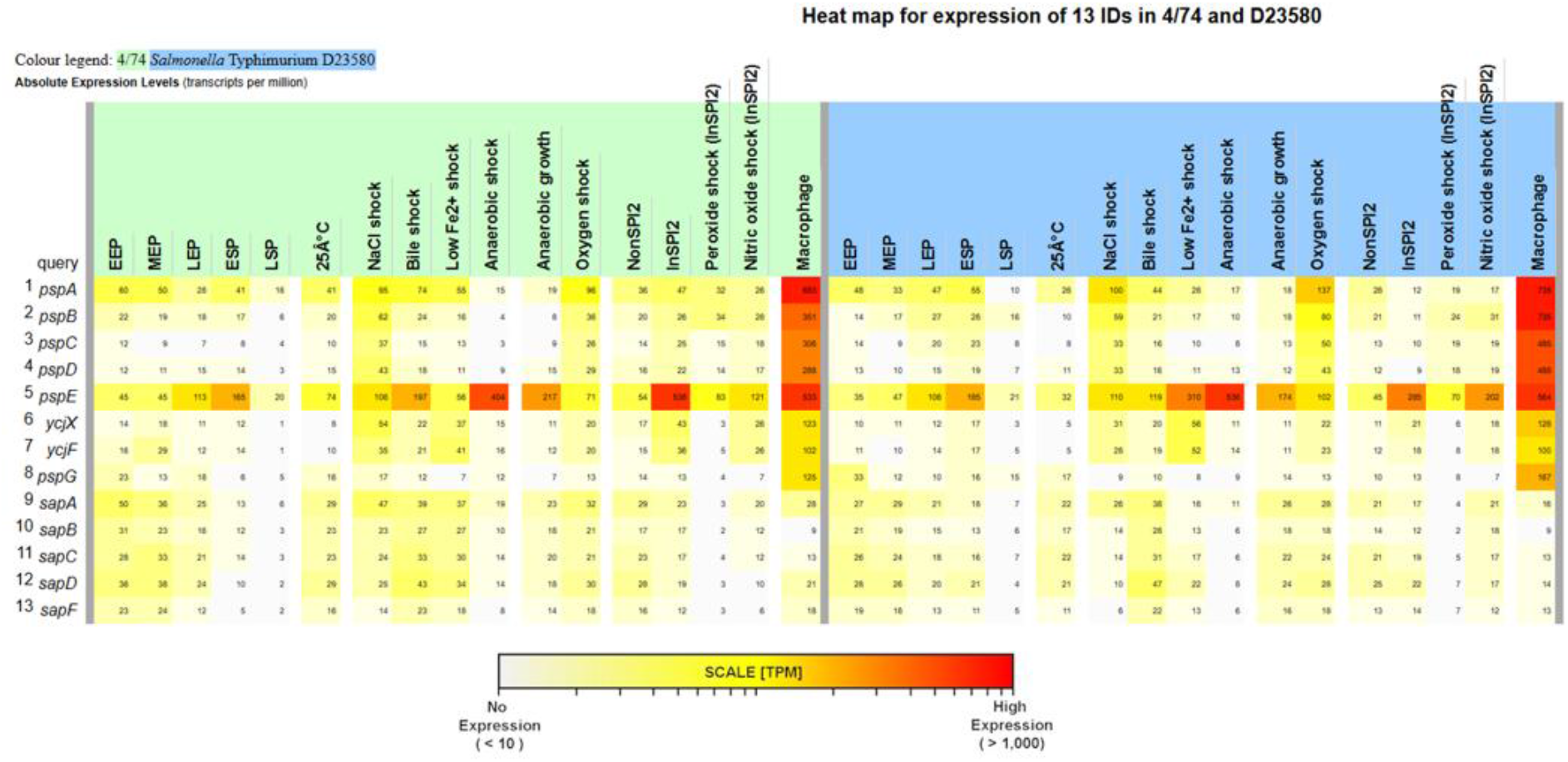
Transcriptomics data shows no evidence of co-regulation between Psp cluster and sap operon in S. Typhimurium. Transcriptomics data for the Psp cluster (pspABCDE-ycjXF and pspG) and sapABCDF operon in S. Typhimurium 4/74 (green) and D23580 (blue). Transcriptomics data has been reproduced from SalComD23580 (https://bioinf.gen.tcd.ie/cgi-bin/salcom_v2.pl?_HL) with the permission of Prof. Jay Hinton (37).

## ABBREVIATIONS

Bac: Biofilm-associated cluster
BLAST: Basic Local Alignment Search Tool
DUF: Domain of Unknown Function
IM30: Inner-membrane-associated 30 kDa protein
iTOL: Interactive Tree of Life
LUCA: Last Universal Common Ancestor
NCBI: National Center for Biotechnology Information
PMF: Proton Motive Force
PspA: Phage shock protein A

